# Type I Interferon regulates cytokine-delayed neutrophil apoptosis, reactive oxygen species production and chemokine expression via activation of p38 MAPK

**DOI:** 10.1101/2020.06.30.179838

**Authors:** Laurence Glennon-Alty, Robert J Moots, Steven W Edwards, Helen L Wright

**Affiliations:** Institute of Life Course and Medical Sciences, University of Liverpool, Liverpool, Merseyside, United Kingdom; Liverpool Health Partners, University of Liverpool, Liverpool, Merseyside, United Kingdom; Clinical Sciences Centre, University Hospital Aintree, Liverpool, Merseyside, United Kingdom; Institute of Infection, Veterinary and Ecological Sciences, University of Liverpool, Liverpool, Merseyside, United Kingdom

**Keywords:** neutrophil, interferon alpha, apoptosis, p38 MAPK

## Abstract

Interferons (IFNs) are key regulators of a number of inflammatory conditions in which neutrophils play an important role in pathology, such as rheumatoid arthritis (RA) and systemic lupus erythematosus (SLE), where Type I IFNs are implicated in disease pathology. However, IFNs are usually generated *in vivo* together with other cytokines that also have immunoregulatory functions but such interactions are poorly-defined experimentally. We measured the effects of Type-I IFN (IFN*α*), elevated in both RA and SLE, on the functions of healthy neutrophils incubated *in vitro* in the absence and presence of pro-inflammatory cytokines typically elevated in inflammatory diseases (TNF*α*, GM-CSF). IFN*α* alone had no effect on neutrophil apoptosis, however it did abrogate the anti-apoptotic effect of GM-CSF (18h, p< 0.01). The enhanced stabilty of the anti-apoptotic protein Mcl-1 and delayed activation of caspase activation normally regulated by GM-CSF were blocked by IFN*α*: this effect was mediated, in part, by activation of p38 MAPK, increased turnover of the anti-apoptotic protein Mcl-1 and cleavage of caspases. IFN*α* alone also primed ROS production alone and maintained the transient priming effect of TNF for up to 4h: it also down-regulated GM-CSF and TNF*α*-activated expression of CXCL1, CXCL2, CXCL3, CXCL8, CCL3 and CCL4, but in contrast increased the expression of CXCL10. These novel data identify complex regulatory signalling networks in which Type I IFNs profoundly alter the response of neutrophils to inflammatory cytokines. This is likely to have important consequences *in vivo* and may explain the complexity and heterogeneity of inflammatory diseases such as RA, in which multiple cytokine cascades have been activated.

## 1 INTRODUCTION

Interferons (IFNs) are a family of cytokine glycoproteins that regulate the immune response to infection by bacteria, viruses and parasites and are broadly grouped into Type-I, -II and -III, based on their receptor signalling (Platanias, 2005;Li et al., 2009). Type-I IFNs, including IFN*α*, IFN*β*, IFN*ϵ*, IFN*κ* and IFN*ω*, signal through the IFN*α* receptor dimer (IFNAR1 and IFNAR2), whereas Type-II IFN, IFN*γ*, signals via the IFN*γ* receptor dimer (IFNGR1 and IFNGR2) (Platanias, 2005). Type-III IFNs, IFN*λ*1, IFN*λ*2, IFN*λ*3 (also known as IL29, IL28*α* and IL28*β*, respectively) signal via a receptor complex comprising IL10R2 and IFNLR1 (Li et al., 2009). Interferon receptors signal through the Janus Kinase (JAK) family of enzymes (JAK1, JAK2, JAK3, Tyk2), which, when activated, induce phosphorylation and nuclear translocation of signal transducer and activation of transcription (STAT) proteins (Platanias, 2005; Li et al., 2009; O’Shea and Plenge, 2012). Type-I interferons activate heterodimers of JAK1:Tyk2 via activation of the IFN*α* receptor complex leading to activation of STAT1:STAT2 heterodimers. In some instances, Type-I interferons can also activate STAT3, STAT4 and STAT5 (Platanias, 2005). Type-II interferons activate heterodimers of JAK1:JAK2 leading to activation of STAT1 homodimers.

As well as playing key roles in the host response to pathogen-mediated infections, IFNs are also implicated in the pathogenesis of a number of inflammatory disorders, including rheumatoid arthritis (RA), systemic lupus erythematosus (SLE), Sjögrens syndrome and myositis (Wright et al., 2014; Higgs et al., 2012; Peck and Nguyen, 2012). In SLE, Type-I interferon signalling correlates with disease severity, and may reflect activation of plasmacytoid dendritic cells (pDCs) which contribute to auto-antibody formation in this disease (Lichtman et al., 2012; Lande et al., 2011). However, the role of IFNs in RA is less well understood and TNF*α* has generally been considered to be the predominant cytokine that drives disease progression, based on the success of therapeutic TNF*α* blockade (TNFi) (Hyrich et al., 2009, 2006).

However, there are several reports of an interferon gene expression signature in RA peripheral blood (Raterman et al., 2012; Reynier et al., 2011; van Baarsen et al., 2010; Mavragani et al., 2010), and we recently identified a correlation between a Type-I ‘IFN-high’ gene expression signature in peripheral blood neutrophils and good response to TNFi in a cohort of RA patients with severe disease activity (Wright et al., 2014, 2017). As TNF*α* inhibits the activity of pDCs and thereby the production of IFN*α* (Palucka et al., 2005), it appears paradoxical that patients who respond to TNFi therapy have high levels of interferon-regulated gene activity. However, whilst TNF*α* can often be detected at high levels in RA synovial fluid (Wright et al., 2012), concentrations of TNF*α* in RA serum are typically low or even undetectable, even in patients who subsequently respond well to TNFi therapy (Wright et al., 2012, 2011).

The effect of Type-II IFN (IFN*γ*) alone on neutrophils has been relatively well defined, and *in vitro* can enhance expression of the high-affinity cell-surface molecules Fc*γ*RI (CD64) (Quayle et al., 1997) and MHC-II (Cross et al., 2003), both of which can be detected on the surface of *ex vivo* neutrophils isolated from RA synovial fluid (Quayle et al., 1997; Cross et al., 2003). IFN-*γ* also delays neutrophil apoptosis and primes the respiratory burst (Sakamoto et al., 2005; Klebanoff et al., 1992). However, the effect of Type-I IFNs on neutrophil function is less clear, with studies reporting contradictory effects on both apoptosis and reactive oxygen species (ROS) production via the respiratory burst (Sakamoto et al., 2005; Koie et al., 2001; Conde et al., 1994).

The aim of this study was to characterise the functional effects of Type-I interferon (IFN*α*) on healthy neutrophils, in the absence and presence of GM-CSF and TNF*α*, cytokines typically elevated in RA. We demonstrate that co-incubation of healthy neutrophils with IFN*α* profoundly alters their functional responses in ways that have important consequences for understanding neutrophil phenotype in inflammatory disease. These data reveal potential complex regulatory cytokine-signalling networks that control immune function in inflammation and inflammatory disease.

## 2 METHODS

### 2.1 Isolation of neutrophils

Participants gave written informed consent according to the declaration of Helsinki and thics approval for this study was obtained from the University of Liverpool Committee on Research Ethics. Neutrophils (purity > 97%, viability > 98%) were isolated from healthy heparinised peripheral blood using HetaSep (Stem Cell) and Ficoll-Paque (GE Healthcare), and contaminating erythrocytes were removed by hypotonic lysis as described previously (Wright et al., 2013). Freshly-isolated neutrophils were suspended at 10^6^ or 5×10^6^ cells/mL in RPMI media (containing 10% human AB serum (Sigma)) and incubated at 37°C and 5% CO_2_ for up to 18h. Cytokines were added at the following concentrations: GM-CSF (5 ng/mL, Roche), TNF (10 ng/mL, Calbiochem), IFN*α*A2 (0.1-20 ng/mL, Sigma), IFN*β*1a (0.1-20 ng/mL, ImmunoTools). Preliminary experiments showed similar results using both Type-I IFNs (IFN*α*A2 and IFN*β*1a), therefore IFN*α*A2 was used throughout and is referred to in the text as IFN*α*. Inhibitors of cell signalling were added 30 min prior to cytokine stimulation: BIRB796 (10*µ*M, p38 MAPK inhibitor, Tocris), PD98059 (50*µ*M, ERK1/2 inhibitor, Merck), JNK-IN-8 (1*µ*M, JNK inhibitor, Selleck Chemicals). Cycloheximide (inhibitor of protein synthesis) was used at 10*µ*g/mL (Sigma) and added 30 min prior to cytokine treatment of neutrophils.

### 2.2 Measurement of apoptosis

Neutrophils (2.5×10^4^) were diluted in 50*µ*L of HBSS (Gibco) containing 0.5*µ*L Annexin V-FITC (Invitrogen), and incubated in the dark at room temperature for 15 min. The total volume was then made up to 250*µ*L with HBSS containing propidium-iodide (PI, 1*µ*g/mL) before analysis by flow cytometry (5,000 events analysed) using a Guava EasyCyte instrument.

### 2.3 qPCR

RNA was extracted using Trizol (Life Technologies) and chloroform, precipitated in isopropanol and cleaned using the RNeasy kit (Qiagen) including a DNase digestion step. cDNA was synthesised using the Superscript III First Strand cDNA Synthesis kit (Invitrogen) using equal concentrations of RNA across samples, as per the manufacturer’s instructions. Real-time PCR analysis was carried out using the QuantiTect SYBR Green PCR kit (Qiagen) as per the manufacturer’s instructions. Analysis was carried out on a Roche 480 LightCycler in a 96-well plate using a 20*µ*L reaction volume. Primers used are shown in Table 1.

**Table 1.**
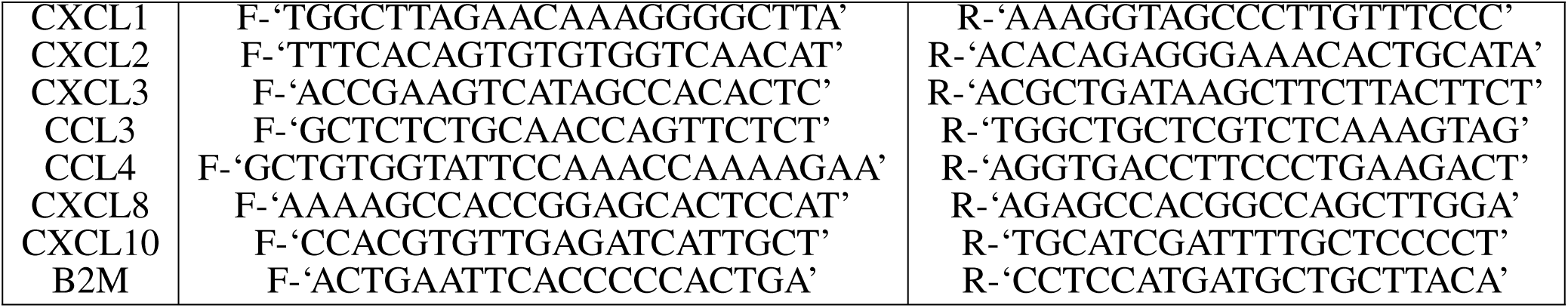
Primers used in qPCR experiments.

Target gene expression was quantified against B2M as a housekeeping gene (Zhang et al., 2005; Pfaffl, 2001). We also carried out an RNA-seq experiment (n=1) to identify targets for validation by qPCR and Western blot. This was carried out using standard RNA-seq protocols on the Illumina HiSeq 2000 platform as previously described (Wright et al., 2017, 2013, 2015).

### 2.4 Western blotting

Neutrophils were rapidly lysed in boiling Laemmli buffer containing phosphatase inhibitors (Calbiochem). Proteins were separated by SDS-PAGE on a 12% gel and transferred onto PVDF membrane (Millipore). Primary antibodies were: Mcl-1 (1:1000, Cell Signalling), caspase-8 (1:1000, Cell Signaling) caspase-9 (1:1000, Cell Signaling), phosphorylated p38 MAPK (1:1000, Cell Signaling), total p38 MAPK (1:1000, Cell Signaling) Actin (1:10,000, Abcam). Secondary antibodies were anti-rabbit IgG (GE Healthcare) and anti-mouse IgG (Sigma) HRP-linked antibodies (1:10,000). Bound antibodies were detected using the ECL system (Millipore) on carefully-exposed film to avoid saturation (Amersham).

### 2.5 Measurement of respiratory burst

Neutrophils (5×10^6^/mL) were incubated for up to 4h with IFN*α* (10 ng/mL) in the absence or presence of GM-CSF (5 ng/mL), or TNF*α* (10 ng/mL). Cells (2×10^5^) were diluted in HBSS containing luminol (10*µ*M) and the respiratory burst stimulated with f-Met-Leu-Phe (fMLP) (1*µ*M). Luminescence was measured, in duplicate, continuously for 30 min.

### 2.6 Statistical analysis

Statistical analysis was carried out using SPSS v20, using the Student’s t-test unless otherwise stated.

## 3 RESULTS

### 3.1 Interferon alpha abrogates GM-CSF delayed neutrophil apoptosis via p38 MAPK signalling

Blood neutrophils constitutively undergo apoptosis after around 24h but during inflammation, apoptosis can be delayed by cytokines such as TNF*α* and GM-CSF (Wright et al., 2013). In order to determine the effect of IFN*α* on neutrophil apoptosis, healthy neutrophils were incubated for 18h in the absence or presence of IFN*α* over a range of concentrations. IFN*α* alone did not significantly alter the rate of constitutive neutrophil apoptosis (Figure 1A), but it dose-dependently abrogated GM-CSF-delayed apoptosis (Figure 1A, p< 0.05). However, this effect was cytokine-specific as it had no significant effect on the rate of apoptosis in neutrophils treated for 18h with TNF*α* (Figure 1B). In order to explore the mechanisms responsible for the effect of IFN*α* on GM-CSF-delayed apoptosis, neutrophils were incubated for up to 18h with these cytokines and protein lysates were analysed by Western blot. GM-CSF has previously been shown to delay neutrophil apoptosis through stabilisation of the anti-apoptotic protein Mcl-1 (Derouet et al., 2004). We found that the levels of Mcl-1 after 6h incubation were significantly lower in neutrophils treated with GM-CSF and IFN*α* compared to GM-CSF alone (Figure 1C,D, p< 0.05). IFN*α* treatment alone had no significant effect on Mcl-1 levels compared to untreated controls (data not shown). Overnight incubations (18h) of neutrophils treated with GM-CSF and IFN*α* had significantly lower levels of pro-caspase-8 and pro-caspase-9 in the presence of IFN*α* compared to GM-CSF alone (Figure 1E-G, p< 0.05).

**Figure 1.**
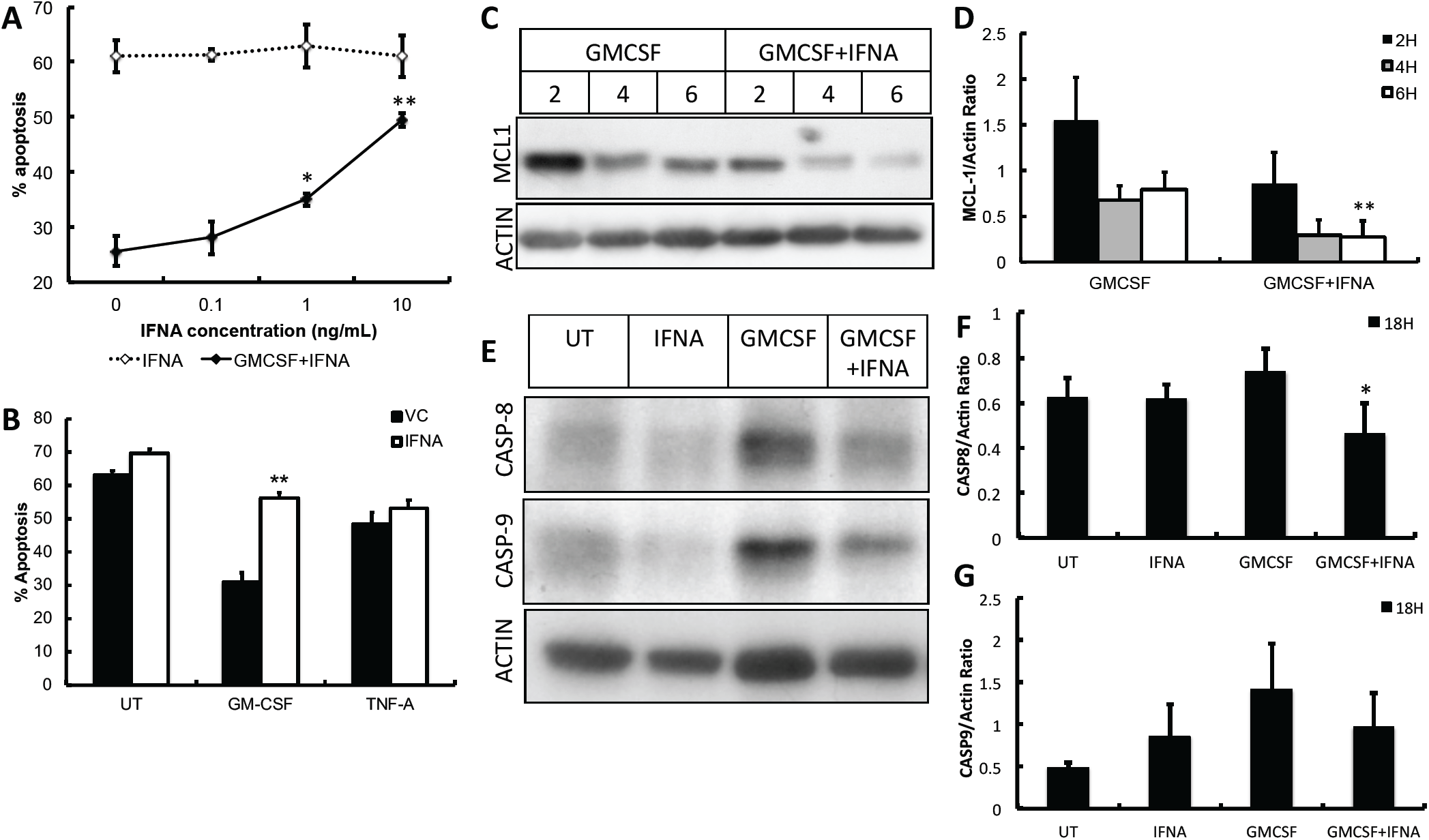
The effect of IFN*α* on neutrophil apoptosis. (A) IFN*α* (IFNA) alone had no effect on neutrophil apoptosis over a range of physiologically-relevant concentrations (0.1-20 ng/mL) compared to untreated neutrophils (VC, vehicle control) at 18h. However, IFN*α* abrogated the anti-apoptotic effect of GM-CSF (5ng/mL). (B) IFN*α* had no effect on the anti-apoptotic effect of TNF*α* (TNF). (C,D) Expression of Mcl-1 was lower in neutrophils co-incubated with GM-CSF and IFN*α* compared to GM-CSF alone. (E-G) The levels of pro-caspase-8 and -9 were lower after 18h culture in the presence of GM-CSF and IFN*α* compared to GM-CSF alone. (all experiments n=3, *p< 0.05, **p< 0.01).

Mcl-1 is an anti-apoptotic, Bcl-2 family protein and a key regulator of neutrophil apoptosis (Moulding et al., 1998; Wardle et al., 2011). It is rapidly turned over in human neutrophils and cellular expression of Mcl-1 correlates with the level of neutrophil apoptosis. However, Mcl-1 levels can be maintained (and apoptosis delayed) either by phosphorylation and stabilisation of existing Mcl-1 protein or synthesis of new Mcl-1 protein (Derouet et al., 2004). Pre-incubation of neutrophils with cycloheximide (CHX), an inhibitor of protein synthesis, prior to addition of GM-CSF and IFN*α* showed that levels of Mcl-1 protein decreased more rapidly in the presence of GM-CSF and IFN*α* compared to GM-CSF alone (Figure 2A). This suggests that IFN*α* blocks the stabilisation of Mcl-1, likely via interference with post-translational modification (phosphorylation) of mature Mcl-1 protein (Thomas et al., 2010). IFN*α* alone had no effect on Mcl-1 levels in the presence of CHX (Figure 2A).

**Figure 2.**
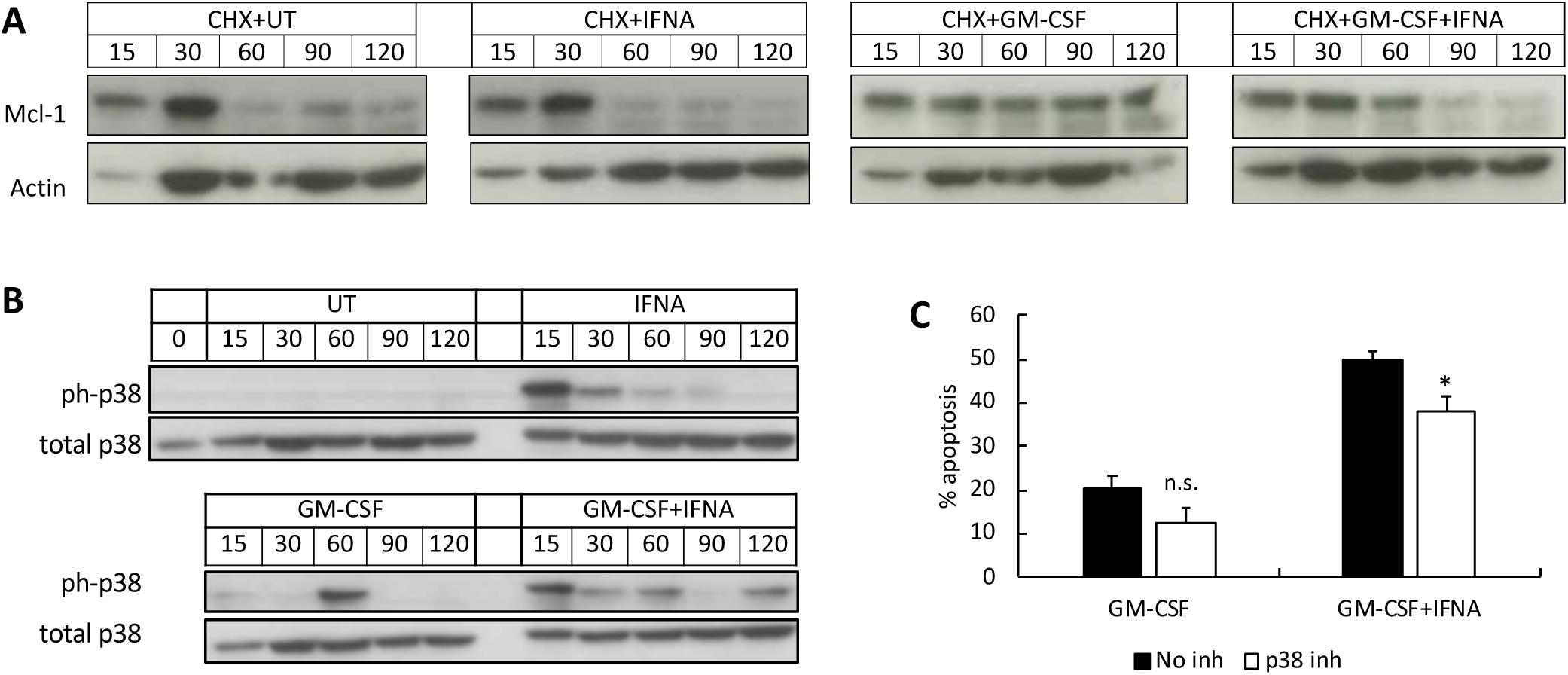
The effect of IFN*α* on Mcl-1 turnover and p38 MAPK activation. (A) Pre-incubation of neutrophils with cycloheximide (CHX) prior to GM-CSF and IFN*α* showed increased turnover of Mcl-1 protein levels in the presence of IFN*α*, likely through blockade of Mcl-1 protein stabilisation. (B) p38 MAPK was rapidly phosphorylated by IFN*α* over 120 min in the absence and presence of GM-CSF. (C) Inhibition of p38 MAPK phosphorylation by BIRB796 significantly inhibited the abrogative effect of IFN*α* on GM-CSF-delayed neutrophil apoptosis over 18h (n=4, *p< 0.05, n.s. = not significant).

We next investigated the potential mechanism by which IFN*α* abrogates the anti-apoptotic effect of GM-CSF using signalling inhibitors of p38 MAPK, JNK and ERK1/2 which are known down-stream targets of the IFNAR1/2 complex (Platanias, 2005). Whilst JNK and ERK1/2 were rapidly activated by IFN*α* in the absence and presence of GM-CSF, inhibition of JNK and ERK1/2 (using JNK-IN-8 and PD98059 respectively) had no significant effect on neutrophil apoptosis (data not shown). However, IFN*α* significantly altered the phosphorylation kinetics of p38 MAPK (Figure 2B), and inhibition of all isoforms of p38 MAPK by BIRB796 significantly decreased the abrogative effect of IFN*α* on GM-CSF-delayed apoptosis by around 25% (Figure 2C, p< 0.05).

### 3.2 Interferon alpha primes the neutrophil respiratory burst and enhances TNF-priming

The production of reactive oxygen species (ROS) via the respiratory burst is rapidly triggered by the phagocytosis of bacteria (Wright et al., 2010). However, in inflammatory diseases such as RA, inappropriate secretion of ROS along with proteolytic enzymes can induce damage to the surface of the joints and degrade cartilage (Wright et al., 2014, 2010). Using luminol-enhanced chemiluminescence, we found that neutrophil ROS production in response to fMLP was not primed by 1h treatment with IFN*α* (Figure 3A). However, ROS production was significantly increased by pre-treatment with IFN*α* for 3h and then stimulated by fMLP (Figure 3B). Co-incubation of IFN*α* with GM-CSF had no significant effect on the GM-CSF-primed respiratory burst, which was sustained over a 5h incubation period (4h timepoint shown in Figure 3C). TNF*α* rapidly primed the neutrophil respiratory burst over short incubation times of up to 1h (data not shown), but by 4h this priming effect was lost. However, co-incubation with IFN*α* significantly sustained this transient priming effect of TNF*α* over and above the priming effect of the IFN*α* alone (Figure 3D).

**Figure 3.**
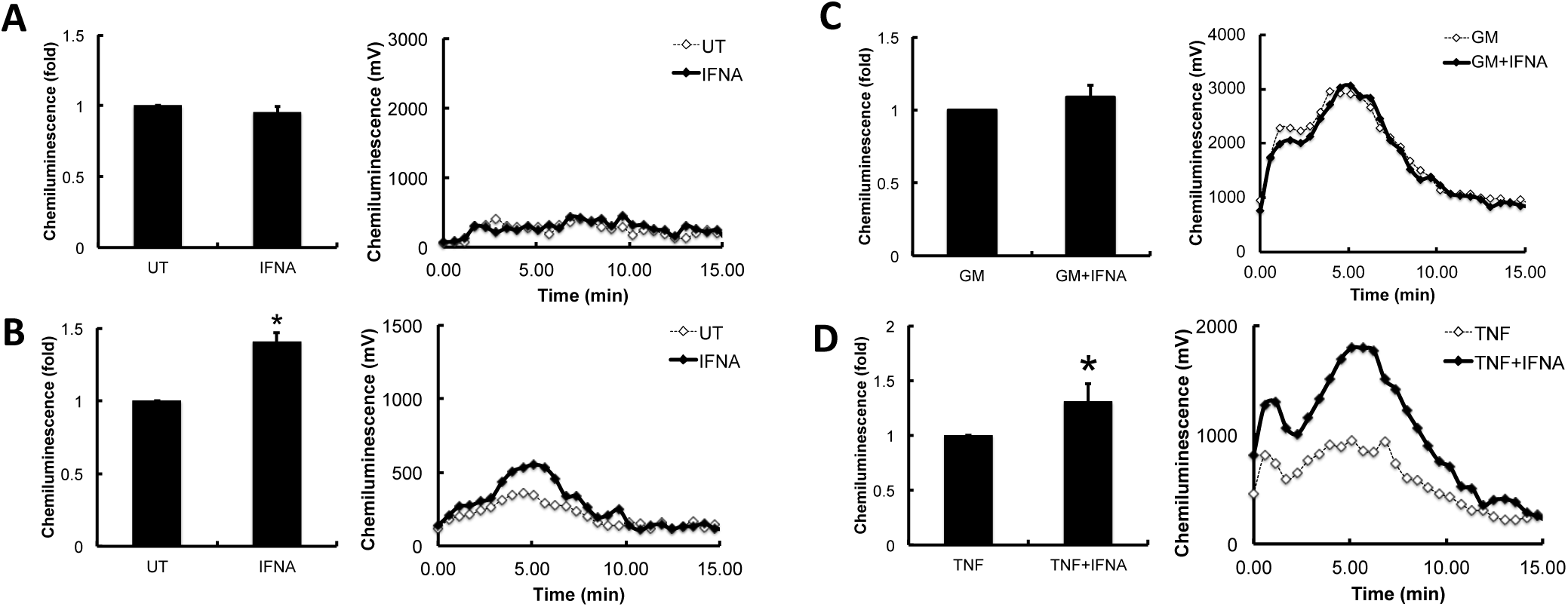
Effect of IFN*α* on neutrophil respiratory burst. (A) IFN*α* (IFNA) did not prime the respiratory burst in response to fMLP (1*µ*M) after short incubation times of 1h, (B) but did significantly prime the respiratory burst after 3h incubation. (C) Co-incubation of IFN*α* with GM-CSF had no effect on neutrophil priming for up to 4h. (D) IFN*α* maintained TNF-induced priming for up to 4h incubation. Representative traces from n=4 experiments (*p< 0.05, **p< 0.01).

### 3.3 Interferon alpha is a key regulator of neutrophil chemokine production

Neutrophils play an important role in the crosstalk between the innate and adaptive immune system via the secretion of chemokines (Mantovani et al., 2011). We measured the expression of chemokines by healthy neutrophils in response to incubation for 1h with GM-CSF and TNF*α* in the absence and presence of IFN*α* using RNA-Seq (n=1, data not shown) and from these data, we selected a panel of genes that were subsequently analysed by qPCR (n=3). GM-CSF and TNF*α* significantly up-regulated the expression of CCL3, CCL4, CXCL1, CXCL2, CXCL3 and CXCL8, but not CXCL10 (Figure 4A,B). IFN*α* alone significantly up-regulated the expression of CXCL10 compared to untreated neutrophils (> 600-fold, data not shown). However, when neutrophils were co-incubated with IFN*α* and GM-CSF or TNF, the increase in expression of CCL3, CCL4, CXCL1, CXCL2, CXCL3 and CXCL8 was significantly abrogated (Figure 4C,D). Conversely, the expression of CXCL10 was significantly enhanced (Figure 4C,D).

**Figure 4.**
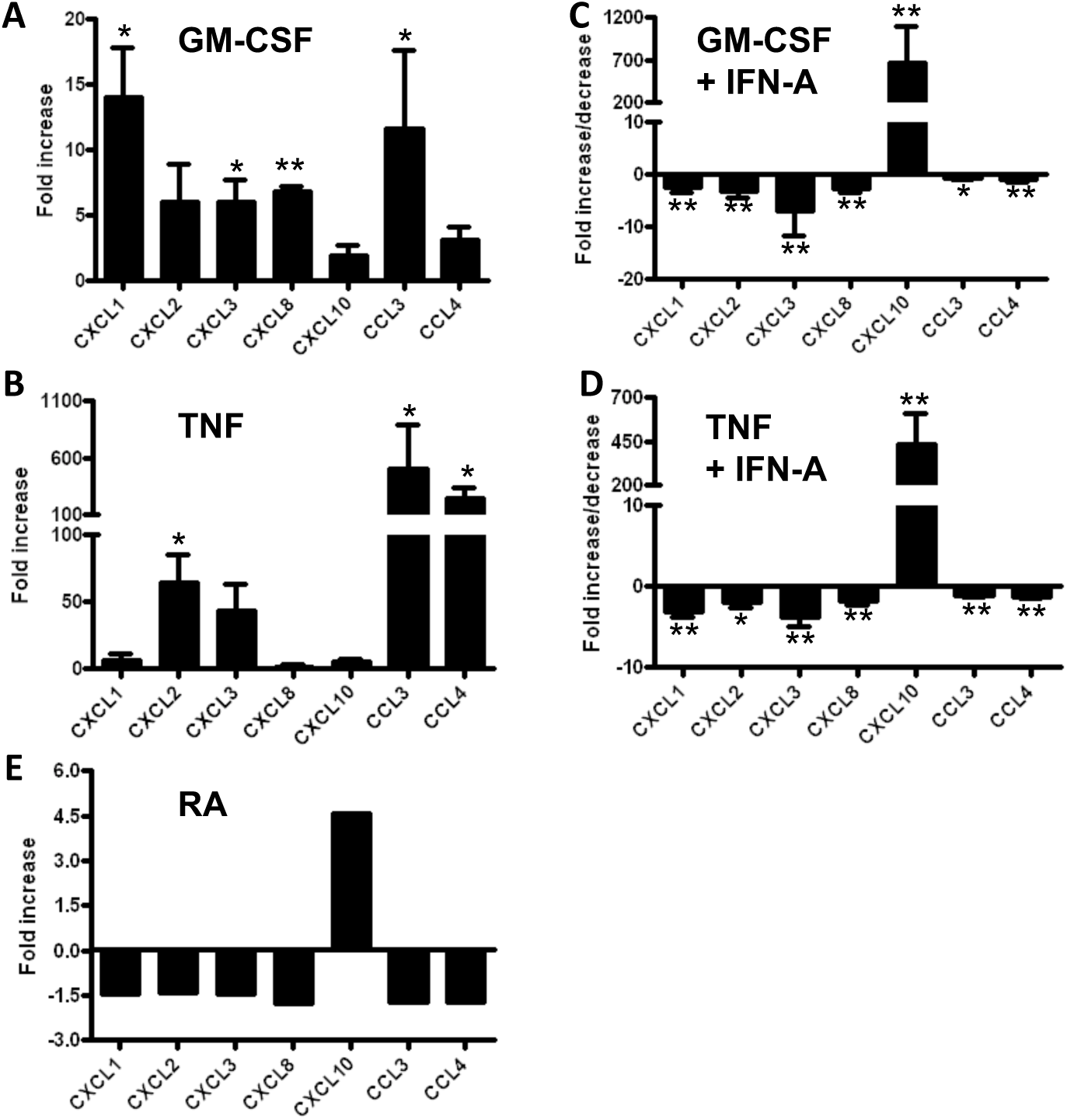
Effect of IFN*α* on chemokine expression. Chemokine expression was upregulated in neutrophils incubated for 1h with (A) GM-CSF or (B) TNF*α* (fold change in expression from 0h shown, n=3 experiments). Co-incubation with IFN*α* (IFN-A) abrogated the expression of chemokines in the presence of (C) GM-CSF or (D) TNF, whereas expression of CXCL10 was enhanced (*p< 0.05, **p< 0.01). (E) RA patients with an ‘IFN-high’ gene expression profile expressed chemokines (CXCL1, CXCL2, CXCL3, CXCL8, CCL3, CCL4) at lower levels than ‘IFN-low’ RA patients (n=14) levels (fold difference in mean expression level between the two groups of patients shown). The expression level of CXCL10 was higher in ‘IFN-high’ RA patients than in ‘IFN-low’ patients.

We next investigated chemokine expression in a group of RA patients who had undergone whole transcriptome sequencing as part of a previous study (Wright et al., 2014). In this study, patients clustered into two groups with either ‘IFN-high’ or ‘IFN-low’ gene expression profiles. When we analysed gene expression levels for CCL3, CCL4, CXCL1, CXCL2, CXCL3 and CXCL8 we found that these were lower in the ‘IFN-high’ patients than in the ‘IFN-low’ patients (Figure 4E), mirroring the effect we observed using qPCR of neutrophils treated *in vitro* with IFNs in combination with TNF*α* and GM-CSF. Interestingly, the expression of CXCL10 was higher in the ‘IFN-high’ patients than in the ‘IFN-low’ patients, confirming that the *in vivo* phenotype of IFN-exposed neutrophils correlates to that induced by *in vitro* exposure to IFNs. These data imply that the *in vitro* regulatory effects of IFNs on neutrophil function that we have observed in this study, may also occur *in vivo*.

## 4 DISCUSSION

While the vast majority of *in vitro* experiments characterise the effects of single pro-inflammatory cytokines on immune cells, it is evident that during inflammation *in vivo*, multiple cytokines are involved in the regulation of immune function. In this study, we have investigated the effect of Type-I interferon (IFN*α*) on neutrophil functions in the absence and presence of two pro-inflammatory cytokines (GM-CSF, TNF*α*) that are implicated in the pathogenesis of rheumatoid arthritis (RA) (Wright et al., 2012). We have demonstrated that IFN*α* profoundly alters the response of neutrophils to these pro-inflammatory cytokines in ways that have important implications for understanding neutrophil phenotype in disease.

In our experiments, IFN*α* abrogated the anti-apoptotic effect of GM-CSF, via blocking the stabilisation of Mcl-1 protein and activation of caspases. We found that IFN*α* altered the phosphorylation kinetics of p38 MAPK, and that blockade of p38 MAPK with a highly selective inhibitor (BIRB796) significantly decreased the effect of IFN*α* on GM-CSF-delayed apoptosis. p38 MAPK is a known regulator of Mcl-1 turnover in neutrophils (Derouet et al., 2004), and inhibition of p38 MAPK using BIRB796 has previously been shown to inhibit Mcl-1 degradation and induction of apoptosis in myeloid cells induced by purvanalol A, another molecule that can abrogate the anti-apoptotic effect of GM-CSF (Phoomvuthisarn et al., 2018). Sodium salicylate also abrogates GM-CSF-delayed neutrophil apoptosis via p38 MAPK signalling and Mcl-1 turnover mechanisms (Derouet et al., 2004). Mcl-1 is an anti-apoptotic Bcl-2 family protein that is a key regulator of neutrophil survival. Activators of p38 MAPK signalling, such as vitamin D, have been shown to have a pro-apoptotic effect on neutrophils via regulation of expression of Bcl-2 and caspase family member genes (Tang et al., 2018; Yang et al., 2015). Whilst we saw changes in the levels of Mcl-1, caspase-8 and caspase-9 protein in our experiments, we did not see any change in expression of these genes in our RNA-seq experiment (data not shown), suggesting the effect of IFN*α* is post-translational. p38 MAPK has also been shown to be anti-apoptotic in neutrophils via the regulation of caspase phosphorylation and activation (Alvarado-Kristensson et al., 2004; Aoshiba et al., 1999) and therefore there remain subtleties in the IFN*α* activation of p38 MAPK that have not been completely addressed in our study. It is interesting to note that IFN*α* did not alter the delay in apoptosis induced by TNF*α* in our study. TNF*α*-delayed apoptosis in human neutrophils is not dependent upon Mcl-1 levels or stabilisation of Mcl-1 protein; this is mediated via NF-*κ*B activation and synthesis of the Bcl-2 family protein A1/Bfl-1 (Cross et al., 2008).

IFN*γ* has previously been shown to prime the neutrophil respiratory burst, but the role of IFN*α* in this process is less clear, with conflicting reports in the literature of the effect of IFN*α* on neutrophil apoptosis and ROS production (Sakamoto et al., 2005; Koie et al., 2001; Conde et al., 1994). This may be due to the IFN*α* isoform that was used in *in vitro* experiments. IFN*α* has 13 different isoforms (Gibbert et al., 2013), and the isoform used in our experiments was IFN*α*A2, which is the most abundant isoform in humans. In addition, inconsistencies in the measurement of IFNs as units/mL rather than ng/mL, often makes direct comparison of experimental conditions difficult. We comprehensively showed that over incubations of up to 1h IFN*α* was not able to prime the respiratory burst. However, over longer incubations > 3h, IFN*α* primed the respiratory burst, both alone and in combination with TNF*α*. This may indicate a requirement for gene expression and a secondary, autocrine signalling effect to fully initiate the primed response.

Our experiments also identified an important role for IFN*α* in the regulation of chemokine production by neutrophils. We showed that IFN*α* significantly decreased the expression of chemokine genes CXCL1, CXCL2, CXCL3, CCL3 and CCL4 in response to GM-CSF and TNF*α*, but enhanced the expression of CXCL10. This has important implications for neutrophil-mediated regulation of the immune response *in vivo*. The chemokines CXCL1, CXCL2 and CXCL3 (GRO-*α*, -*β* and –*γ*, respectively) and CCL3 and CCL4 (MIP-1*α* and -1*β*, respectively) predominantly function as neutrophil chemoattractants, whereas CXCL10 (IP-10) is primarily a chemoattractant for T-cells, monocytes, NK cells and dendritic cells. Therefore, the cellular signalling events regulated by combinations of inflammatory cytokines (GM-CSF and/or TNF*α*) and IFNs *in vivo* may act as a switch to direct the immune response away from neutrophil-mediated inflammation towards lymphocyte-mediated inflammation, underlying the key role of neutrophils in the cross-talk between the innate and adaptive immune systems (Mantovani et al., 2011). Importantly, we noted that the changes in chemokine expression in the presence of IFN*α* mirrored exactly the phenotype that is seen in RA neutrophils, underlying the important role of IFN*α* in the regulation and fine-tuning of neutrophil function in inflammatory disease.

In summary, our data show that the response of human neutrophils to TNF*α* and GM-CSF is profoundly altered *in vitro* by IFN*α*, whereby IFN*α* acts as a switch to alter the inflammatory neutrophil phenotype by regulating apoptosis, ROS release and chemokine production This has important consequences for neutrophil inflammatory responses *in vivo*, particularly in complex and heterogeneous inflammatory diseases such as RA in which combinations of these cytokines exist, and may explain why some patients respond better to certain biologic therapies than others.

## AUTHOR CONTRIBUTIONS

LGA carried out the experiments and analysed the data. HLW designed the research, carried out the experiments, analysed the data, and wrote the manuscript. RJM revised the manuscript. SWE analysed the data and revised the manuscript.

## ACKNOWLEDGMENTS

None.

## REFERENCES

Alvarado-Kristensson, M., Melander, F., Leandersson, K., Ronnstrand, L., Wernstedt, C., and Andersson, T. (2004). p38-mapk signals survival by phosphorylation of caspase-8 and caspase-3 in human neutrophils. J Exp Med 199, 449–58. doi:10.1084/jem.20031771

Aoshiba, K., Yasui, S., Hayashi, M., Tamaoki, J., and Nagai, A. (1999). Role of p38-mitogen-activated protein kinase in spontaneous apoptosis of human neutrophils. J Immunol 162, 1692–700

Conde, M., Andrade, J., Bedoya, F. J., and Sobrino, F. (1994). Inhibitory effect of interferon-alpha on respiratory burst and glucose metabolism in phagocytic cells. J Interferon Res 14, 11–6

Cross, A., Bucknall, R. C., Cassatella, M. A., Edwards, S. W., and Moots, R. J. (2003). Synovial fluid neutrophils transcribe and express class ii major histocompatibility complex molecules in rheumatoid arthritis. Arthritis Rheum 48, 2796–806

Cross, A., Moots, R. J., and Edwards, S. W. (2008). The dual effects of tnfalpha on neutrophil apoptosis are mediated via differential effects on expression of mcl-1 and bfl-1. Blood 111, 878–84

Derouet, M., Thomas, L., Cross, A., Moots, R. J., and Edwards, S. W. (2004). Granulocyte macrophage colony-stimulating factor signaling and proteasome inhibition delay neutrophil apoptosis by increasing the stability of mcl-1. J Biol Chem 279, 26915–21. doi:10.1074/jbc.M313875200

Gibbert, K., Schlaak, J. F., Yang, D., and Dittmer, U. (2013). Ifn-alpha subtypes: distinct biological activities in anti-viral therapy. Br J Pharmacol 168, 1048–58. doi:10.1111/bph.12010

Higgs, B. W., Zhu, W., Richman, L., Fiorentino, D. F., Greenberg, S. A., Jallal, B., et al. (2012). Identification of activated cytokine pathways in the blood of systemic lupus erythematosus, myositis, rheumatoid arthritis, and scleroderma patients. Int J Rheum Dis 15, 25–35. doi:10.1111/j.1756-185X.2011.01654.x

Hyrich, K. L., Deighton, C., Watson, K. D., Consortium, B. C. C., Symmons, D. P., Lunt, M., et al. (2009). Benefit of anti-tnf therapy in rheumatoid arthritis patients with moderate disease activity. Rheumatology (Oxford) 48, 1323–7. doi:10.1093/rheumatology/kep242

Hyrich, K. L., Watson, K. D., Silman, A., Symmons, D. P., and British Society for Rheumatology Biologics, R. (2006). Predictors of response to anti-tnf-alpha therapy among patients with rheumatoid arthritis: results from the british society for rheumatology biologics register. Rheumatology (Oxford) 45, 1558–65. doi:10.1093/rheumatology/kel149

Klebanoff, S. J., Olszowski, S., Van Voorhis, W. C., Ledbetter, J. A., Waltersdorph, A. M., and Schlechte, K. G. (1992). Effects of gamma-interferon on human neutrophils: protection from deterioration on storage. Blood 80, 225–34

Koie, T., Suzuki, K., Shimoyama, T., Umeda, T., Nakaji, S., and Sugawara, K. (2001). Effect of interferon-alpha on production of reactive oxygen species by human neutrophils. Luminescence 16, 39–43. doi:10.1002/bio.606

Lande, R., Ganguly, D., Facchinetti, V., Frasca, L., Conrad, C., Gregorio, J., et al. (2011). Neutrophils activate plasmacytoid dendritic cells by releasing self-dna-peptide complexes in systemic lupus erythematosus. Sci Transl Med 3, 73ra19. doi:10.1126/scitranslmed.3001180

Li, M., Liu, X., Zhou, Y., and Su, S. B. (2009). Interferon-lambdas: the modulators of antivirus, antitumor, and immune responses. J Leukoc Biol 86, 23–32. doi:10.1189/jlb.1208761

Lichtman, E. I., Helfgott, S. M., and Kriegel, M. A. (2012). Emerging therapies for systemic lupus erythematosus–focus on targeting interferon-alpha. Clin Immunol 143, 210–21. doi:10.1016/j.clim.2012.03.005

Mantovani, A., Cassatella, M. A., Costantini, C., and Jaillon, S. (2011). Neutrophils in the activation and regulation of innate and adaptive immunity. Nat Rev Immunol 11, 519–31. doi:10.1038/nri3024

Mavragani, C. P., La, D. T., Stohl, W., and Crow, M. K. (2010). Association of the response to tumor necrosis factor antagonists with plasma type i interferon activity and interferon-beta/alpha ratios in rheumatoid arthritis patients: a post hoc analysis of a predominantly hispanic cohort. Arthritis Rheum 62, 392–401. doi:10.1002/art.27226

Moulding, D. A., Quayle, J. A., Hart, C. A., and Edwards, S. W. (1998). Mcl-1 expression in human neutrophils: regulation by cytokines and correlation with cell survival. Blood 92, 2495–502

O’Shea, J. J. and Plenge, R. (2012). Jak and stat signaling molecules in immunoregulation and immune-mediated disease. Immunity 36, 542–50. doi:10.1016/j.immuni.2012.03.014

Palucka, A. K., Blanck, J. P., Bennett, L., Pascual, V., and Banchereau, J. (2005). Cross-regulation of tnf and ifn-alpha in autoimmune diseases. Proc Natl Acad Sci 102, 3372–7. doi:10.1073/pnas.0408506102

Peck, A. B. and Nguyen, C. Q. (2012). Transcriptome analysis of the interferon-signature defining the autoimmune process of sjogren’s syndrome. Scand J Immunol 76, 237–45. doi:10.1111/j.1365-3083.2012.02749.x

Pfaffl, M. W. (2001). A new mathematical model for relative quantification in real-time rt-pcr. Nucleic Acids Research 29, 2002–2007

Phoomvuthisarn, P., Cross, A., Glennon-Alty, L., Wright, H. L., and Edwards, S. W. (2018). The cdk inhibitor purvalanol a induces neutrophil apoptosis and increases the turnover rate of mcl-1: potential role of p38-mapk in regulation of mcl-1 turnover. Clin Exp Immunol 192, 171–180. doi:10.1111/cei.13107

Platanias, L. C. (2005). Mechanisms of type-i- and type-ii-interferon-mediated signalling. Nat Rev Immunol 5, 375–86. doi:10.1038/nri1604

Quayle, J. A., Watson, F., Bucknall, R. C., and Edwards, S. W. (1997). Neutrophils from the synovial fluid of patients with rheumatoid arthritis express the high affinity immunoglobulin g receptor, fc gamma ri (cd64): role of immune complexes and cytokines in induction of receptor expression. Immunology 91, 266–73

Raterman, H. G., Vosslamber, S., de Ridder, S., Nurmohamed, M. T., Lems, W. F., Boers, M., et al. (2012). The interferon type i signature towards prediction of non-response to rituximab in rheumatoid arthritis patients. Arthritis Res Ther 14, R95. doi:ar3819[pii]10.1186/ar3819

Reynier, F., Petit, F., Paye, M., Turrel-Davin, F., Imbert, P. E., Hot, A., et al. (2011). Importance of correlation between gene expression levels: application to the type i interferon signature in rheumatoid arthritis. PLoS One 6, e24828. doi:10.1371/journal.pone.0024828PONE-D-11-05657[pii]

Sakamoto, E., Hato, F., Kato, T., Sakamoto, C., Akahori, M., Hino, M., et al. (2005). Type i and type ii interferons delay human neutrophil apoptosis via activation of stat3 and up-regulation of cellular inhibitor of apoptosis 2. J Leukoc Biol 78, 301–9. doi:10.1189/jlb.1104690

Tang, Y., Liu, J., Yan, Y., Fang, H., Guo, C., Xie, R., et al. (2018). 1,25-dihydroxyvitamin-d3 promotes neutrophil apoptosis in periodontitis with type 2 diabetes mellitus patients via the p38/mapk pathway. Medicine (Baltimore) 97, e13903. doi:10.1097/MD.0000000000013903

Thomas, L. W., Lam, C., and Edwards, S. W. (2010). Mcl-1; the molecular regulation of protein function. FEBS Lett 584, 2981–9. doi:S0014-5793(10)00465-5[pii]10.1016/j.febslet.2010.05.061

van Baarsen, L. G., Wijbrandts, C. A., Rustenburg, F., Cantaert, T., van der Pouw Kraan, T. C.1, Baeten, D. L., et al. (2010). Regulation of ifn response gene activity during infliximab treatment in rheumatoid arthritis is associated with clinical response to treatment. Arthritis Res Ther 12, R11. doi:ar2912[pii]10.1186/ar2912

Wardle, D. J., Burgon, J., Sabroe, I., Bingle, C. D., Whyte, M. K., and Renshaw, S. A. (2011). Effective caspase inhibition blocks neutrophil apoptosis and reveals mcl-1 as both a regulator and a target of neutrophil caspase activation. PLoS One 6, e15768. doi:10.1371/journal.pone.0015768

Wright, H. L., Bucknall, R. C., Moots, R. J., and Edwards, S. W. (2012). Analysis of sf and plasma cytokines provides insights into the mechanisms of inflammatory arthritis and may predict response to therapy. Rheumatology 51, 451–9. doi:10.1093/rheumatology/ker338

Wright, H. L., Chikura, B., Bucknall, R. C., Moots, R. J., and Edwards, S. W. (2011). Changes in expression of membrane tnf, nf-kappab activation and neutrophil apoptosis during active and resolved inflammation. Ann Rheum Dis 70, 537–43. doi:10.1136/ard.2010.138065

Wright, H. L., Cox, T., Moots, R. J., and Edwards, S. W. (2017). Neutrophil biomarkers predict response to therapy with tumor necrosis factor inhibitors in rheumatoid arthritis. J Leukoc Biol 101, 785–795. doi:10.1189/jlb.5A0616-258R

Wright, H. L., Moots, R. J., Bucknall, R. C., and Edwards, S. W. (2010). Neutrophil function in inflammation and inflammatory diseases. Rheumatology 49, 1618–31. doi:10.1093/rheumatology/keq045

Wright, H. L., Moots, R. J., and Edwards, S. W. (2014). The multifactorial role of neutrophils in rheumatoid arthritis. Nat Rev Rheumatol 10, 593–601. doi:10.1038/nrrheum.2014.80

Wright, H. L., Thomas, H. B., Moots, R. J., and Edwards, S. W. (2013). Rna-seq reveals activation of both common and cytokine-specific pathways following neutrophil priming. PLoS One 8, e58598. doi:PONE-D-12-34202

Wright, H. L., Thomas, H. B., Moots, R. J., and Edwards, S. W. (2015). Interferon gene expression signature in rheumatoid arthritis neutrophils correlates with a good response to tnfi therapy. Rheumatology (Oxford) 54, 188–93. doi:10.1093/rheumatology/keu299

Yang, H., Long, F., Zhang, Y., Yu, R., Zhang, P., Li, W., et al. (2015). 1alpha,25-dihydroxyvitamin d3 induces neutrophil apoptosis through the p38 mapk signaling pathway in chronic obstructive pulmonary disease patients. PLoS One 10, e0120515. doi:10.1371/journal.pone.0120515

Zhang, X., Ding, L., and Sandford, A. J. (2005). Selection of reference genes for gene expression studies in human neutrophils by real-time pcr. BMC Mol Biol 6, 4. doi:10.1186/1471-2199-6-4

